# Feedback control of neurogenesis by tissue packing

**DOI:** 10.1101/252445

**Authors:** Tom W. Hiscock, Joel B. Miesfeld, Kishore R. Mosaliganti, Brian A. Link, Sean G. Megason

## Abstract

Balancing the rate of differentiation and proliferation in developing tissues is essential to produce organs of robust size and composition. Whilst many molecular regulators have been established, how these connect to physical and geometrical aspects of tissue architecture is poorly understood. Here, using high-resolution timelapse imaging, we find that dense tissue packing and complex cell geometries play a significant role in regulating differentiation rate in the zebrafish neural tube. Specifically, in regions of high cell density, progenitors are physically pushed away from the apical surface, which, in a Notch-dependent manner, leads to their differentiation. Using simulations we show that this naturally performs negative feedback control on cell number. Our results suggest a model whereby differentiation rate is carefully tuned to correct fluctuations in cell number, originating from variable cell cycle progression and inherently probabilistic differentiation programs.

## Introduction

Growth is a central process in developmental programs, and its control is critical to generate tissues of a particular size. In many tissues, this growth is substantial. For example, cell number in the retina increases ∼400-fold during its development (Alexiades and Cepko, 1996), but occurs with a highly stereotyped rate and duration to ensure that the final number of cells in the tissue is tightly controlled.

Tissue growth rate, here defined as the rate of increase in cell number within the tissue, affects two interlinked and essential processes of development: (1) proliferation of a pool of dividing progenitors, which increases progenitor number, and (2) differentiation of progenitors into post-mitotic differentiated cells, which reduces progenitor number (with progenitor apoptosis typically negligible). In homeostatic tissues, proliferation and differentiation must be perfectly balanced to maintain a constant pool of cycling cells. However, in developing tissues, proliferation and differentiation must instead be coordinated (Hardwick and Philpott, 2014; Hindley and Philpott, 2012; Kicheva et al., 2014), so that early, proliferation dominates and the tissue grows, whereas late, differentiation increases relative to proliferation and the growth rate finally approaches zero (Miguez, 2015). It is key to know how these two processes are tuned as development progresses in order to understand how a tissue reaches a specified final size. Furthermore, stereotyped tissue growth must occur despite large variability in proliferation rates (i.e. cell cycle times), and inherently probabilistic modes of differentiation (He et al., 2012). Determining how differentiation and proliferation are controlled – in the engineering sense of correcting errors – is thus fundamental to understanding how tissues reach a *robust* final size, despite the stochastic and noisy mechanisms underpinning their development.

Here, we focus on the neural tube as a model system of growth control, which shows stereotypic growth dynamics - specifically an initial phase of rapid proliferation, followed by a gradual shift to differentiation (Saade et al., 2013) Much is known about the molecular regulators of neural tube growth. For example, the hes/her transcription factors promote and maintain proliferation of the progenitor pool, whereas expression of genes such as neurogenin or p27 induces cell cycle exit and differentiation (reviewed in (Hardwick et al., 2015). Differentiation is also affected by the inheritance of specific protein domains following division (Alexandre et al., 2010; Dong et al., 2012; Huttner and Kosodo, 2005; Noctor et al., 2004; Paolini et al., 2015). Expression and genetic analyses also implicate a number of extracellular regulators of cell fate. Some of these are local signals, such as the Delta-Notch pathway, whereas others, such as Wnt, TGFbeta, Shh and BMP, can act over a longer range (Dessaud et al., 2007; Garcia-Campmany and Marti, 2007; Le Dreau et al., 2014; Saade et al., 2013; Zechner et al., 2003). These then define molecular gradients that give rise to differential differentiation rates across the dorsal-ventral axis of the neural tube (Kicheva et al., 2014).

We wondered whether there are also mechanical, or physical, regulators of proliferation and/or differentiation during neural tube growth. Experiments on cell stretching *in vitro* reveal that proliferation can respond significantly to externally applied mechanical strain (Aragona et al., 2013; Benham-Pyle et al., 2015; Streichan et al., 2014). Furthermore, differentiation of various stem cells in culture has been shown to be highly dependent on the mechanical properties of their microenvironment (Arulmoli et al., 2015; Engler et al., 2006; Gilbert et al., 2010; Leipzig and Shoichet, 2009; Pan et al., 2016; Seidlits et al., 2010). However, the extent to which these observations apply to neural tube development is unclear, particularly since it has a much more complex tissue architecture than the 2D cell monolayers that have been studied previously. This tissue architecture is: (1) epithelial, and therefore has a distinct apical-basal polarity; (2) pseudostratified, in which multiple nuclei are densely packed at different depths within a single epithelial layer; and (3) highly dynamic, with apical mitoses driving extensive rearrangement of nuclei, termed “interkinetic nuclear migration” (Leung et al., 2011; Norden et al., 2009). To what extent these properties impact differentiation in this tissue is unknown.

Here we uncover a significant role for the physical and geometric aspect of tissue packing during neural tube development. Using high resolution *in toto* timelapse imaging (Megason, 2009), we show that crowding at the apical surface leads to an increased rate of differentiation within the tissue. At the single cell level, this manifests itself as a correlation between cells whose nuclei have been displaced basally (due to apical crowding) and those that differentiate. We then show, using simulations, that such a feedback can naturally guide robust developmental trajectories in the face of probabilistic differentiation processes and highly variable cell cycle progression. Given the prevalence of similar pseudostratified tissue architectures, both in developmental contexts, (e.g. cortex (Kosodo et al., 2011), retina (Leung et al., 2011), pancreas (Bort et al., 2006)), as well as in homeostatic adult tissues (e.g. the intestine (Grosse et al., 2011; Jinguji and Ishikawa, 1992)), we speculate that tissue packing and apical crowding may be a widely-used regulator of differentiation and growth across a range of different organisms and tissues.

## Results

### The neural tube is densely-packed and crowded at the apical surface

To investigate neurogenesis in the zebrafish neural tube, we collected high resolution confocal stacks of embryos doubly transgenic for a ubiquitous membrane label *Tg(actb2:memCherry2)* (Xiong et al., 2014), and a pan-neuronal marker *Tg(neurod:eGFP)* (Obholzer et al., 2008), one of the earliest markers of neural differentiation (Lee, 1997) (Fig. 1A, Fig. S1C). For measurement we define differentiation based on expression of *neurod* rather than cell cycle exit. Our tracking data suggests these are tightly correlated since we did not observe *neurod* in dividing cells (0/91).

**Figure 1:**
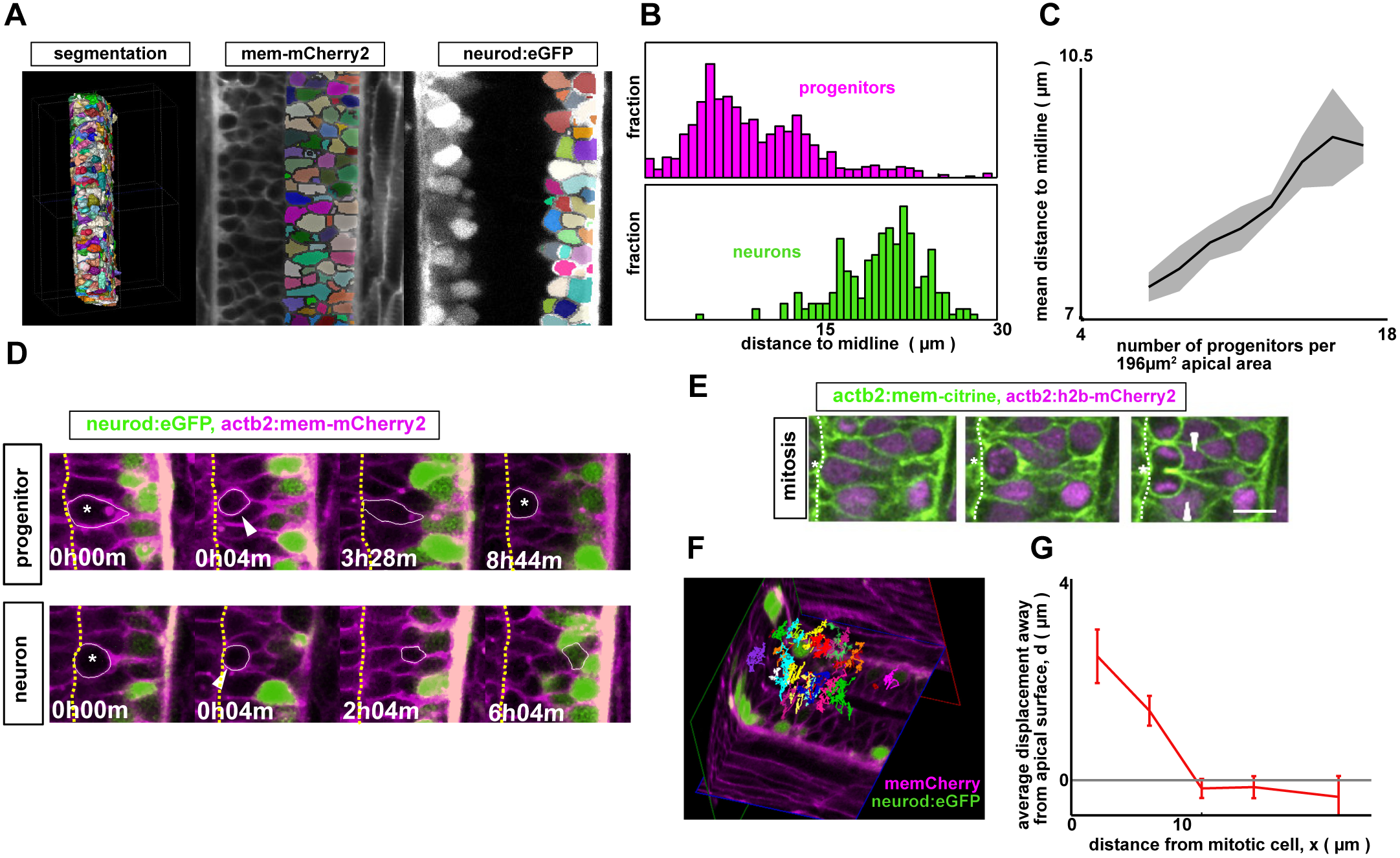
The neural tube is a densely packed and dynamic pseudostratified epithelium. A: Analysis pipeline: membrane-labeled images are segmented and cropped using custom scripts to generate 3D cell meshes for the entire neural tube. Each mesh is then classified as a neuron or a progenitor, according to the expression level of the neural marker (*neurod:eGFP*) (See also Fig. S1C). B: Distance of cell centroid position to the midline, for both progenitors and neurons. C: Regions of high progenitor density correlate with regions where progenitors are located far from the midline (shown is mean distance plus/minus standard error). D: Some representative cell tracks from Go Figure2. Upper: a cycling progenitor. Lower: a nascent neuron. Asterisks denote a mitotic cell; arrowheads denote one of the daughter cells from the mitotic cell. E: Endogenous mitotic cells physically deform their neighbors, depicted in a montage of images separated by 9 minutes per frame. Asterisk denotes mitotic cell ending cytokinesis; arrowheads denote perturbed adjacent cells. (Scale bar: 10*μ*m) F: Cell tracking from high resolution, *in toto* timelapse movies is performed in GoFigure2, reveals extensive nuclear movement. G: Mitotic cells transiently push their neighbors away from the apical surface. We measure the maximal basal displacement moved by a given cell as a result of a nearby division. We also record the distance x *μ*m along the apical surface between the measured cell and the mitotic cell. We then group the data according to x, and plot the mean and standard error as shown.

3D cell segmentations were generated using ACME (Mosaliganti et al., 2012) and revealed a densely-packed, pseudostratified epithelial tissue architecture (Fig. 1A). Consistent with other neuroepithelia, neurons are located basally, whereas progenitors are predominantly apical (Fig. 1B), although remain attached to both the apical and basal surface (Fig. 2A). However, there is a large variability in the distance of the progenitor nucleus (approximated by the cell segmentation centroid, Fig. S1) to the midline. This is typical of pseudostratified epithelia in which there are multiple nuclei at different distances to the midline within a densely packed single cell layer.

**Figure 2:**
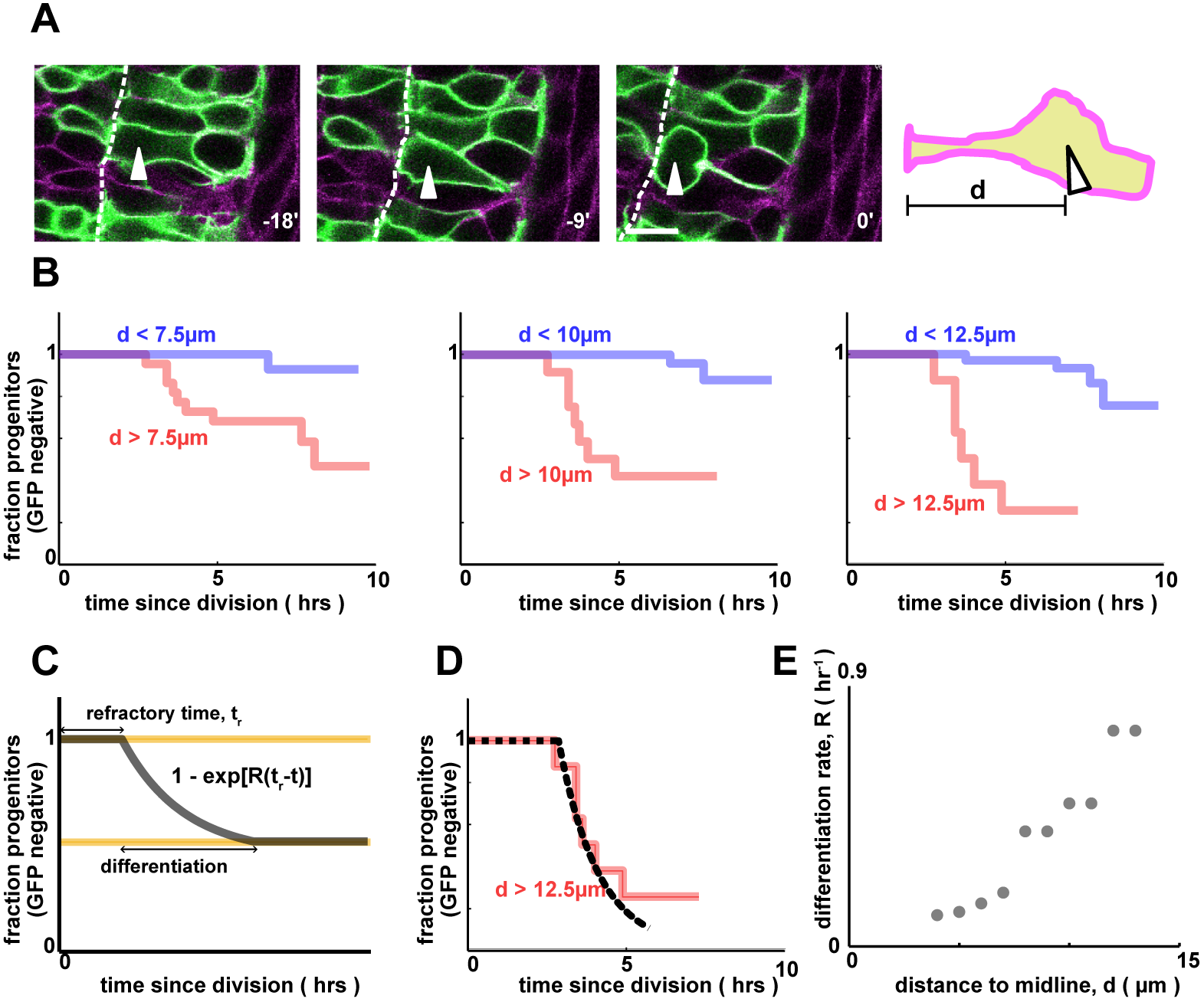
Progenitors that are far from the apical surface differentiate more frequently. A: Quantification of pre-mitotic cell shape by distance to midline, *d*. (Scale bar: 10*μ*m) B: Single cell tracking reveals that cells that are far from the apical surface predivision turn on *neurod* more rapidly than those that are close. The dependence of differentiation rate on cell shape is independent of the threshold value that defines which cells are ‘far’ and which cells are ‘close’ (Figure 6.4). (Left: p = 0.01; middle: p = 1e-6; right: p = 1e-7; n = 58). C: A simple model for differentiation, where for some time window, each cell differentiates at a constant rate per unit time, *R*, which we call the differentiation rate. D: Numerical fit of the model in (C) to the data in (B). E: Differentiation rate, *R,* as a function of the distance to midline, *d*.

We hypothesized that the variability in nuclear position was reflective of variability in progenitor number at different positions within the neural tube. Indeed we find that the density of progenitors (per unit apical surface) varies across the tissue, as does the density of neurons. However, we see a clear correlation between progenitor density and nuclear position. Specifically, in regions where there are many progenitors per unit apical surface, their mean distance to the midline is higher (Fig. 1C). This follows from a purely geometric argument: more progenitors produces crowding at the apical surface, thereby forcing some progenitor nuclei to be displaced basally. In this way, there is a direct geometrical connection between epithelial cell density and the distribution of nuclear depths due to cell packing.

Next, we collected *in toto* timelapse imaging datasets that allowed single cell tracking of neural progenitors over ∼12hrs of development starting from 24hpf (Xiong et al., 2013). These data revealed the highly dynamic aspect of tissue architecture, as evidenced by the significant movement seen in tracking single nuclei over time (Fig. 1F). By following individual progenitors, we see substantial, but largely undirected movement between divisions. As progenitors differentiate, they move basally (Fig. 1D), and as they divide, they move apically (Fig. 1E). A consequence of this is that the surrounding cells are significantly deformed, moving their nuclei away from the apical surface (Fig. 1G). Therefore, similar to the retina, an increase in pressure at the apical surface caused by mitotic cells drive substantial movement of nuclei within the tissue (known as interkinetic nuclear migration) (Leung et al., 2011).

### Progenitors far from the apical surface differentiate

Next, we used the timelapse data as a sensitive, single cell assay to measure differentiation rates, by directly tracking progenitors and assigning them to a neural identity based on *neurod:eGFP* expression. To avoid biases caused by variations along the DV axis, we restricted analysis to cells located within the central ∼30% of the neural tube. By collecting many such tracks, we could generate Kaplan Meier plots (commonly known as ‘survival curves’ in the medical literature), as shown in Fig. 2C that characterize the rate at which progenitors differentiate (Rich et al., 2010). Kaplan-Meier curves are insensitive to incomplete cell tracks, avoids effects of cells moving out of frame, or the timelapse ending before a cell has definitively divided or differentiated.

Unexpectedly, we saw a correlation between the differentiation of progenitors and their geometry. To quantify this, we analyzed the shapes of progenitors prior to their division. Restricting our shape measurements to the pre-mitotic mother cell was key in order to say something about causation, since it is known that progenitors undergo stereotyped cell shape changes as they differentiate, which would result in a trivial correlation between cell shape and differentiation. As a simple measure of cell shape, we measured the maximum distance of the cell nucleus to the apical surface observed in a time window 45-60 minutes prior to mitosis (Fig. 2A). We explicitly ignored any transient basal movement induced by neighboring cells as they divided, as we hypothesized that the long-term cell shape would be more informative (in Fig. S2, we confirm that the transient displacements have minimal effects on differentiation).

We then asked if this pre-mitotic shape correlated with *neurod* expression dynamics in the daughter cells. Strikingly, we found a strong association between the cells whose nuclei were far from the apical surface, and the cells that rapidly turned on *neurod* after dividing (Fig. 2B). By fitting a simple parametric form to the Kaplan Meier differentiation curves (Fig. 2C,D), we could quantify how the differentiation rate, *R*, depended on distance of nucleus to the midline, *d*, and found a significant positive correlation between the two (Fig. 2E). Interestingly, a similar observation has been made in the vertebrate retina (Baye and Link, 2007), suggesting that this could be a rather general feature of neuroepithelia.

Together with the observations of tissue packing in Fig. 1, this suggests a model whereby apical crowding induces differentiation. More specifically, apical pressure – a result of a high density of cells at the apical surface – causes progenitors to be displaced away from the apical surface, which in turn leads to an increase in differentiation rate. Conversely, in regions of low apical pressure (i.e. few cells), we would expect a lower rate of differentiation.

### Pushing progenitor nuclei away from the apical surface by an arrested mitotic cell promotes differentiation

To test this hypothesis, we aimed to locally increase crowding at the apical surface and thereby push progenitors away from the midline. To do this, we exploited the fact that mitotic cells significantly deform their neighbors, a result of their large size and rounded morphology at the apical surface upon division (Fig. 1G). Therefore we could mimic an increase in crowding at the apical surface simply using a mitotic cell that is prevented from dividing. To achieve this, we arrested a small fraction of cells in mitosis, by inducing expression of a dominant negative version of PLK1, a kinase necessary for mitotic exit (Strzyz et al., 2015). Following heatshock-induced mosaic dnPLK1 expression, the small fraction of cells that were expressing the construct (BFP positive) failed to exit mitosis and remained rounded and apical for extended periods of time (Fig. 3A). Further, these arrested mitotic cells substantially deformed the shapes of neighboring progenitors and, as hoped, caused a significant increase in the distance of cell nuclei to the apical surface (Fig. 3B). We then measured whether such a perturbation to apical crowding and cell shape impacted the proliferation and/or differentiation of these cells.

**Figure 3:**
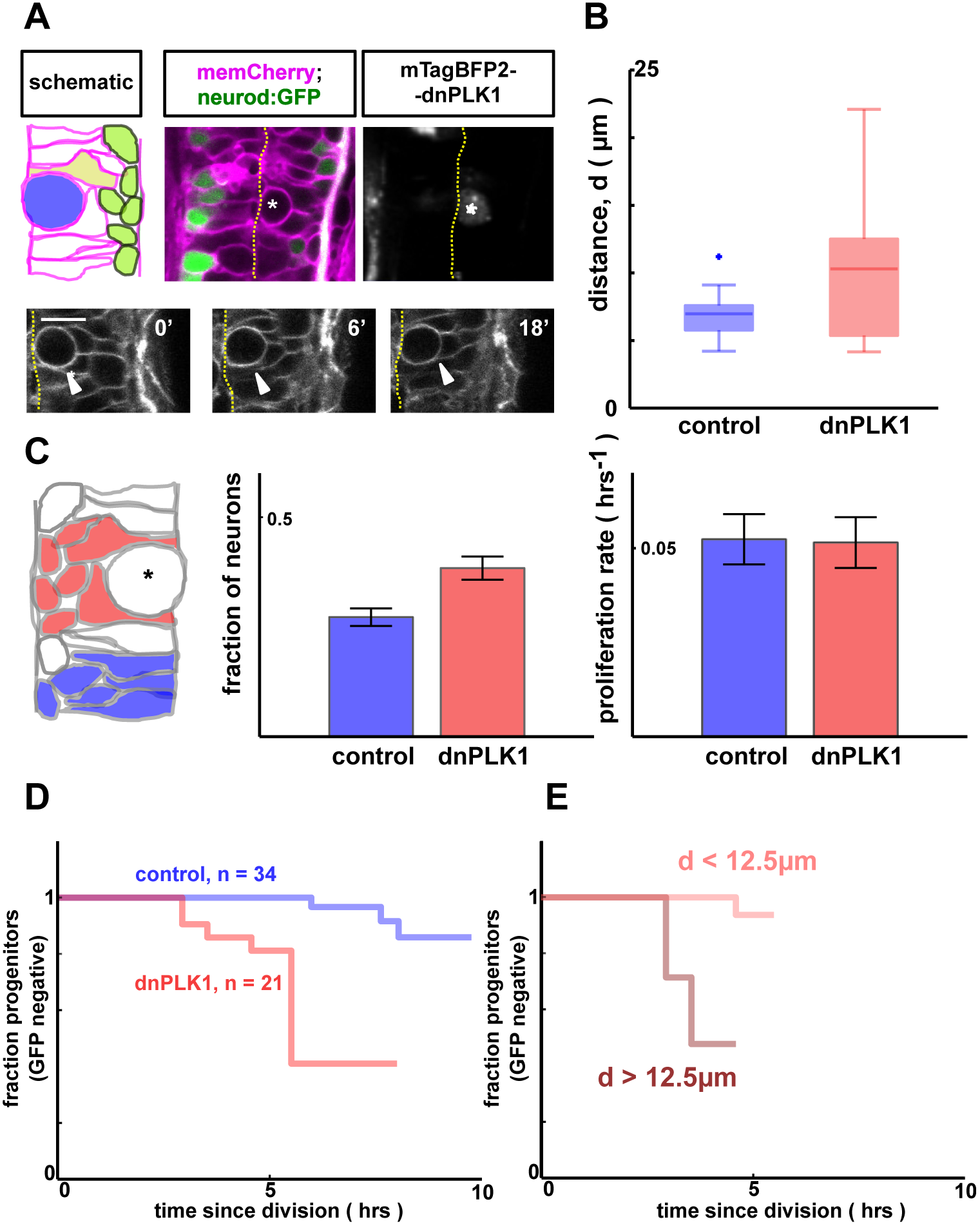
Pushing progenitors away from the apical surface by an arrested mitotic cell promotes their differentiation. (A) Cell shapes are perturbed by inducibly and mosaically arresting neighbouring cells in mitosis, using a heat-shock inducible dnPLK1 construct, which prevents mitotic exit (lower). (Scale bar: 10*μ*m) (B) Cells adjacent to the arrested mitotic cell are shifted basally (p < 0.01, n = 10 for dnPLK1, n = 26 for control). Here, *d* is the distance to the apical surface prior to division, as measured in Fig. 2. (C) There is an increased fraction of neurons adjacent to arrested mitotic cells (red) than in control regions without an arrested cell (blue) (p < 0.01). Proliferation rates are similar in the two cases (right) (p = 0.9). (n = 7 for both cases) (D) Tracking of single cells adjacent to an arrested mitotic cell reveals a significant increase in differentiation (p < 0.01), compared to control (data from the same experiment as Fig. 2). (E) Single cell tracking reveals the same correlation between cell shape and *neurod* dynamics as in Fig. 2 for cells adjacent to arrested mitotic cells (p < 0.001, using the n = 21 tracked dnPLK1 cells).

We used two separate methods to assay differentiation rates. First, we collected high-resolution confocal stacks to count neuron and progenitor numbers following prolonged deformation by arrested mitotic cells. We found that there was a significant increase in the number of neurons in close proximity to an arrested cell, compared to unperturbed control regions from the same embryo, indicating an increase in the differentiation rate (Fig. 3C). We then measured proliferation rates by counting precursors at two time points, and subtracting to get the number of division events (Fig S3B). No significant difference was found in proliferation rate between regions deformed by an arrested cell and unperturbed regions (Fig. 3C). Together, this suggests that crowding progenitor nuclei away from the apical surface leads to an increase in differentiation, but minimal changes to proliferation, consistent with our previous results.

Secondly, we generated *in toto* timelapse datasets of these perturbed embryos and tracked cells that were in close proximity to the mitotically-arrested cells (but were themselves not arrested i.e. BFP negative). Tracking data revealed that progenitors adjacent to the arrested mitotic cell more rapidly and extensively turned on *neurod* expression than in control embryos (Fig. 3D). Furthermore, within this dataset, we saw the same correlation between pre-mitotic cell shape and its daughter’s *neurod* dynamics as above i.e. those progenitors that were pushed far from the apical surface by the arrested cell were exactly those that rapidly turned on *neurod* following division (Fig. 3E).

### The effect of the apical arrested mitotic cell is primarily physical

The strong correlation between cell shape and differentiation rate in response to neighboring mitotic cells suggests that the effect of the arrested cell on its neighbors depends on its ability to physically deform them. However it is conceivable that a non-physical mechanism such as expression of some secreted molecule or cell surface protein by mitotic cells could also affect differentiation rate in neighbors. To test this possibility, we sought to arrest mitotic cells in a way that they did not increase pressure at the apical surface and so does not deform their neighbors to the same extent. Inspired by previous studies on neuroepithelial nuclear migration, we co-expressed p50 in the dnPLK1-arrested cells, which is known to inhibit the dynactin complex and thus impair apical movement of nuclei (Burkhardt et al., 1997; Tsuda et al., 2010; Tsujikawa et al., 2007). Indeed, we found a small number of progenitors that were arrested in mitosis (assayed by condensed chromosomes, Fig. 4A), but were non-apical and therefore did not push neighboring cell nuclei away from the apical surface (Fig. 4B). By tracking cells adjacent to these basal arrested cells, we no longer observed a local increase in differentiation, suggesting that the apical location of the mitotic cell is necessary for its neurogenic effect (Fig. 4C). These results suggest that the extent of crowding at the apical surface, as parameterized by nuclear position, strongly influences the differentiation rate of neural progenitors.

**Figure 4:**
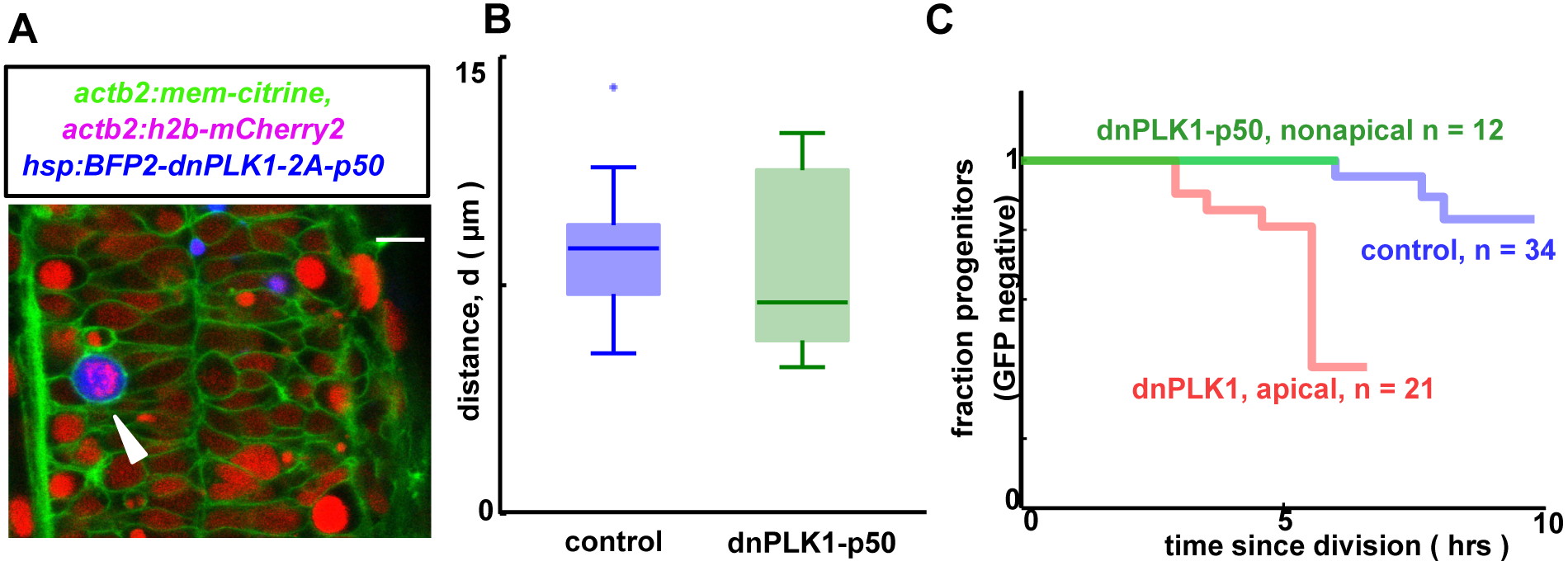
Progenitors adjacent to a non-apical mitotic cell do not differentiate more rapidly relative to control. A: Arrested mitotic cells can be nonapical (arrowhead). h2b-mCherry demonstrates condensed chromosomes hence mitotic entry. Scale bar: 10*μ*m) B: Progenitors are not pushed away from the apical surface adjacent to dnPLK1-p50 arrested non-apical mitotic cells prior to division (p = 0.6) C: Non-apical mitotic cells do not induce differentiation of their neighbors (p = 0.8, n = 12 vs. control, which is the same control data as in Fig. 3).

### Notch as a candidate molecular transducer of apical crowding

Next, we aimed to understand how apical crowding and cell shape could be sensed molecularly, and therefore how the physical effect of tissue packing connects to the molecular circuitry upstream of neural differentiation. We started by considering the Notch pathway (Bray, 2016) whose activity is necessary for progenitor maintenance in the zebafish neural tube. This pathway is particularly appealing since previous studies in the retina have suggested that Notch activity can depend on nuclear positioning via an apical-basal gradient of ligand and receptor (Aggarwal et al., 2016; Clark et al., 2012; Del Bene et al., 2008; Hatakeyama et al., 2014; Latasa et al., 2009).

To test whether Notch was involved in shape-sensing, we measured Notch activity in cells that were significantly and persistently deformed by an adjacent hsp:dnPLK1 arrested mitotic cell . We used a novel transgenic reporter to mark Notch activity, which drives destabilized GFP expression downstream of the *nort* promoter (Fig. S4A), a known direct target of Notch (Tsutsumi and Itoh, 2007). The transgene expressed fluorescence in a manner nearly identical to expression of the endogenous transcript, including robust expression in spinal cord neural progenitors (Fig. 5A, S4B). Importantly, abatement of Notch signaling by knockdown of Rbpj (Fig. S4C) or expression of dominant negative Maml (not shown), essential cofactors for transcriptional activity of all Notch subtypes resulted in nearly complete loss of reporter fluorescence. In addition, over-expression of Notch1 intracellular domain (NICD1) strongly enhanced reporter activity in either endogenous sites of nort expression or within ectopic locations that normally do not express nort (Fig. S4D).

**Figure 5:**
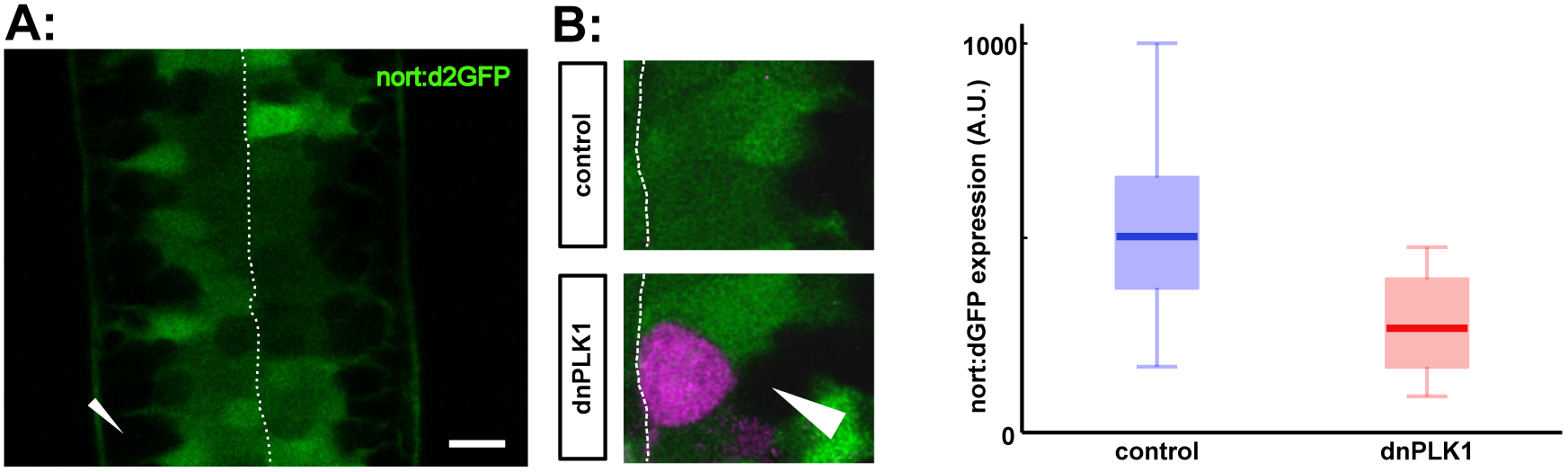
Notch activity as a potential readout of nuclear position. A: nort:d2GFP expression reports Notch activity in the neural tube. Arrorhead denotes GFP-negative, basally localized neurons. (Scale bar: 10*μ*m) B: Notch activity (green) is significantly inhibited in progenitors that are adjacent to an arrested mitotic cell (magenta) (p < 0.01, n = 15 for control, n = 12 for dnPLK1). (Scale bar: 10*μ*m)

We hypothesized that the arrested mitotic cells would locally inhibit Notch activity to drive differentiation. Consistent with this hypothesis, we saw a significant downregulation of Notch activity in progenitors adjacent to an arrested mitotic cell (Fig. 5B). Given that Notch is required for progenitor maintenance in the neural tube (Appel et al., 2001; Huang et al., 2012; Schier et al., 1996; Yeo and Chitnis, 2007), this supports a model whereby apical crowding inhibits Notch signaling, thus causing cells to differentiate. However, the extent to which Notch is sufficient to explain this phenomenon, and the mechanism by which the pathway responds to cell shape, is yet to be determined (see Discussion).

### Apical crowding provides a negative feedback on progenitor number

Regardless of how it is transduced molecularly, the effect of tissue packing and apical crowding on differentiation rate would naturally provide a negative feedback between growth rate and cell number in this tissue, analogous to previous theoretical work on growth control in imaginal discs (Shraiman, 2005). Specifically, as the number of progenitors increases within the tissue, we expect an increase in the pressure and/or crowding of cells at the apical surface. This will then change the distribution of cell shapes within the tissue, giving rise to a higher number of basally located progenitors with smaller apical area, and thus a higher rate of differentiation. Therefore, as the number of progenitors within a region increases, their rate of differentiation also increases. This in turn leads to a depletion of the progenitor pool, giving rise to a negative feedback on progenitor cell number (Fig. 6A).

**Figure 6:**
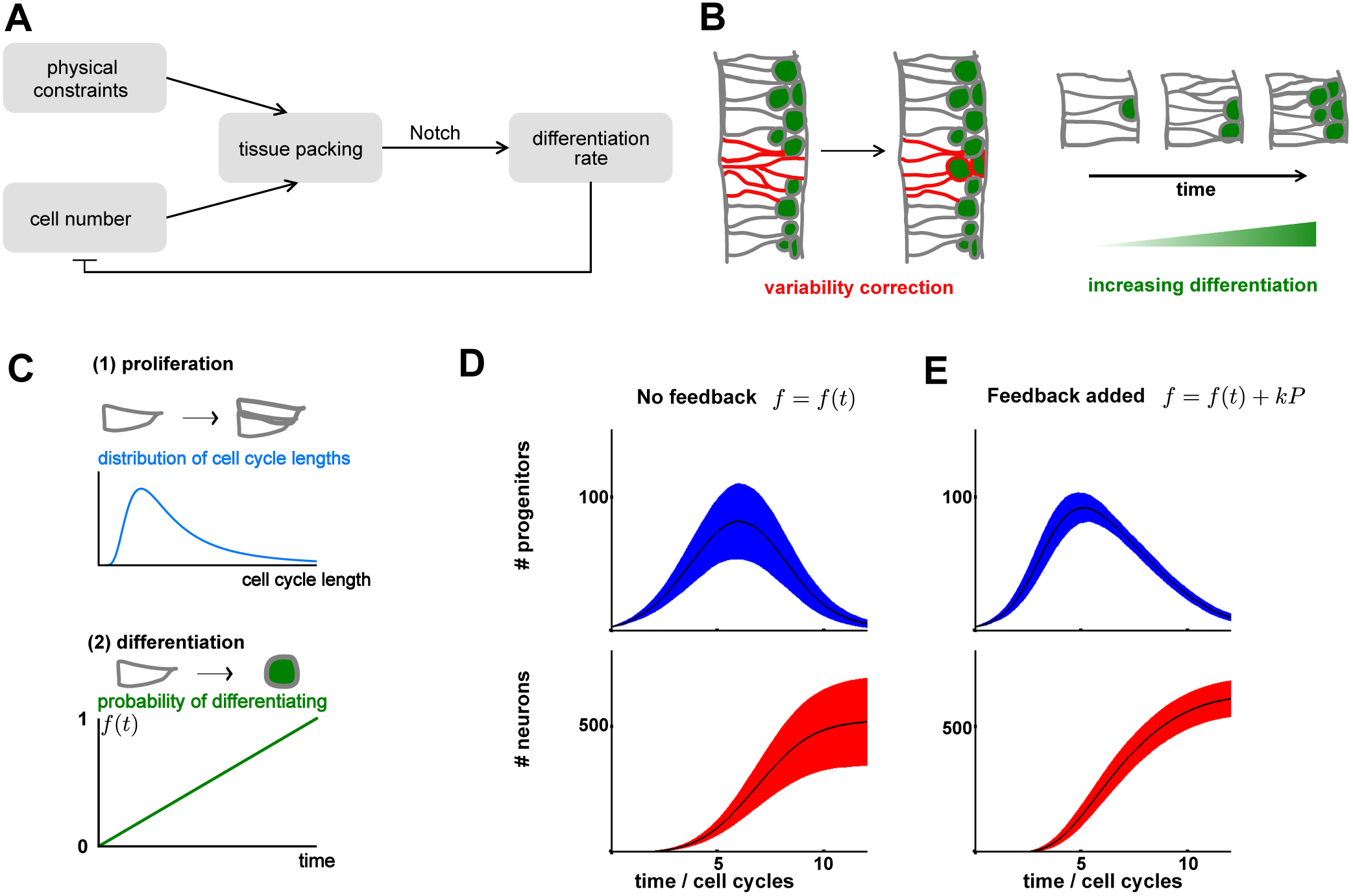
Apical crowding as a feedback mechanism to control progenitor number. A: We hypothesize that the regulation of differentiation by cell shape forms a negative feedback loop. We speculate that this feedback could perform several functions, schematized in B. B: Negative feedback naturally reduces variability in progenitor number. Left: a region of high progenitor density (red) corrects itself by differentiating. Right: If cells divide within a confined space, negative feedback predicts an increase in differentiation over time. C: An *in silico* model of neural tube development with two main ingredients. Upper: progenitors proliferate with a given cell cycle distribution. Lower: progenitors differentiate shortly after dividing, with probability *f(t)*. D: Progenitor and neuron numbers are variable without feedback. The black line is the mean trajectory; solid regions denote mean plus/minus standard deviation for 3000 independent simulations. E: When we add feedback (by allowing *f(t)* = *f*_0_*(t)* + *κP*), the standard deviation in neuron and progenitor number is significantly reduced.

One possible role for this negative feedback would be to suppress fluctuations. Neurogenesis dynamics are far from deterministic, and must operate despite highly variable cell cycle lengths and multiple stochastic influences on the differentiation machinery (He et al., 2012). This generates significant variability in neuron and progenitor numbers across the tissue, and between different embryos. A crucial role for negative feedback would be to reduce this variability: intuitively, regions with too many progenitors would compensate by differentiating more; and regions with too few progenitors by differentiating less (see Fig. 6B).

To formally test this intuition, we constructed a mathematical model of neural tube development. We assume progenitors proliferate with a certain distribution of cell cycle times (Fig. 6C). Then, after dividing, there is a certain probability a given progenitor will either self renew (*f*_*PP*_), asymmetrically divide (*f*_*NP*_) or generate two post-mitotic daughters (*f*_*NN*_). Motivated by work in the chick neural tube (Saade et al., 2013), we assume that each daughter cell differentiates independently with probability *f* (i.e. division mode probabilities *f*_*PP*_ = *f*^2^, *f*_*NP*_ = 2*f*(1-*f*), *f*_*NN*_ = (1-*f*)^2^ form a binomial distribution). Over time, we assume that this differentiation probability increases (i.e. *f(t)* is an increasing function of time, *t*), so that initially the tissue grows, and later on differentiation dominates and the pool of precursors is depleted. We find that, without any feedback mechanism, the numbers of progenitors and neurons are highly variable in the *in silico* neural tube (Fig. 6D), a consequence of the probabilistic differentiation and variable cell cycle lengths. However, if we incorporate the feedback from apical crowding – whereby the differentiation probability, *f*(*t,P*), increases not only as a function of time, but also as a function of progenitor number, *P* – then we find that this variability is significantly reduced, consistent with our intuition.

## Discussion

In this study, using a combination of *in toto* timelapse imaging and physical perturbations, we have identified apical crowding as a novel regulatory mechanism for neurogenesis. In particular, we found that when neural progenitors are squeezed away from the midline (and/or compressed at their apical surface), they were more likely to differentiate. This suggests that neurogenesis dynamics within the neural tube are not entirely deterministic, nor cell-autonomously programmed, and instead can be regulated by the mechanical properties of the tissue, its environment and how these interact to regulate tissue packing. Using modeling, we argued that this phenomenon results in negative feedback between progenitor number and differentiation rate, and that this can significantly reduce variability in developmental trajectories.

The negative feedback module also gives a mechanism to coordinate changes in tissue size and growth rate over developmental time. In particular, during early neural tube development, there are few progenitors and the tissue is relatively loosely packed, and thus in our model differentiation is rather low. However, as the neural tube continues to grow, it becomes compressed by the tissues surrounding it (likely the skin, somites and notochord, which each compress the neural tube from different directions), causing cells to be densely packed and so more likely to differentiate (see Fig. 6B). In this way, the exit from the early proliferative phase of neural tube growth could be governed by this mechanical feedback, in addition to known molecular regulators (Hudish et al., 2016), and therefore growth continues until all the available space is filled. This hypothesis may provide an explanation for the hyperproliferation phenotypes in human open neural tube defects (NTDs), such as spina bifida, in which the spinal cord is ‘open’ or exposed at birth (Copp et al., 2013). We speculate that the increased growth is a result of the reduction in physical constraints acting on the neural tube. This has been directly observed in surgical models of NTDs, in which surgically removing the skin overlying the spinal cord results in increased proliferation in chick embryos (You et al., 1994). However, more experiments are required to determine to what extent such a space-filling mechanism is actually operating in the zebrafish neural tube, and its significance during unperturbed development.

In this work, we have largely focused on explaining our observations at the level of cells and tissues. Preliminary work has implicated the Delta-Notch signaling pathway as a potential mechanism by which cells measure their shape, although the precise details are far from clear. One hypothesis is that, given that Notch ligand and receptor are both apically enriched, one might expect the level of active nuclear Notch (NICD) to depend on the distance of the nucleus to the apical surface, provided NICD is rapidly degraded (or bound by an inhibitor e.g. *numb*) en route to the nucleus (Aggarwal et al., 2016). In this case, having the nucleus in close proximity to the apical Notch receptors gives a higher chance that a given NICD molecule reaches the nucleus and activates transcription.

However, there are other possibilities. In particular, whilst we have focused on nuclear position as a readout of cell shape, there could also be a role for apical contact area. Correspondingly, another possible mechanism is that, as proposed elsewhere (Clark et al., 2012; Shaya et al., 2017), the amount of Notch signaling received by a cell is dependent on the size of its cell-cell contacts with neighboring cells (which is directly related to its apical area), since this is where the bulk of the Notch receptor is located. Therefore, a cell with smaller apical area will have a smaller contact with neighboring Delta positive cells and consequently will receive lower active Notch signaling. A further possibility is that it is not just geometry but also force that is at play. Notch signaling has been shown to depend, at the single molecular level, on forces and therefore the forces associated with apical compression, could be modulating Notch activity directly, rather than indirectly via its effect on cell geometry (Gordon et al., 2015). These possible mechanisms are not mutually exclusive and aspects of each may be coordinated to regulate neurogenesis. Testing these hypotheses will require higher resolution tools to measure and perturb Notch signal transduction.

Whilst in this work we have focused on Notch activity as a readout of cell shape, it is likely that other signaling pathways are involved. The WNT pathway is a promising candidate, since it is known to be responsive to mechanical cues (Brunet et al., 2013; Fernandez-Sanchez et al., 2015; Nowell et al., 2016) and has significant effects on neurogenesis (Zechner et al., 2003). Other mechanotransduction pathways such as the Hippo pathway (Dupont et al., 2011) or the piezo proteins (Coste et al., 2012) could also be determining the response to increased pressure at the apical surface. Finally, it is possible that it is not just apical crowding, but also signals from the basal compartment (e.g. TGFbetas secreted by basally positioned neurons) that is important. Elucidating the molecular details of the shape-based feedback mechanism, and the interactions between Notch and other signaling pathways and the apical surface should be the subject of further work.

Finally, our work may yield important insights to understanding how differentiation and proliferation are balanced more generally. Many tissues have a similar architecture (i.e. densely packed, pseudostratified epithelia, with a large degree of nuclear movement), most notably other neuroepithelia, but also a range of other developmental and adult tissues (Spear and Erickson, 2012). It will be interesting to determine whether the feedback between tissue packing and differentiation described in this work is a common feature in these tissues, and to understand how its deregulation could lead to novel tissue architectures, such as the folded primate brain (Otani et al., 2016; Tallinen et al., 2014), or aberrant growth during tumorigenesis (Fernandez-Sanchez et al., 2015; Ou and Weaver, 2015).

## Materials and Methods

### Zebrafish strains and maintenance

*Tg(neurod:eGFP)* (Obholzer et al., 2008), *Tg(actb2:mem-mCherry2)* (Xiong et al., 2014), Tg(*crystA α*:Gal4) (Hayes et al., 2012) and *Tg(actb2:mem-citrine-citrine)* (Xiong et al., 2013) (referred to as “mem-citrine”) have been described previously. *Tg(actb2:h2b-mCherry2)* was generated using a plasmid that encodes the h2b sequence fused to mCherry2, in a pMTB backbone as described previously (Xiong et al., 2014). *Tg(-3.5kb nort:d2GFP)* was constructed using Gateway^®^ recombination and the Tol2 Kit (Kwan et al., 2007) to place the *3.5kb nort* promoter upstream of d2GFP (destabilized GFP; Clonetech). Natural spawning was used, and embryos were incubated at 28°C throughout their development (including during imaging), but excluding small amounts of time during experimental manipulation (such as microinjection, mounting) which occurred at room temperature.

Zebrafish work was approved by the Harvard Medical Area Standing Committee on Animals under protocol number 04487.

### Confocal imaging

Embryos were anaesthetized in two different ways depending on the type of experiment. First, for continuous timelapse imaging, alpha bungarotoxin was delivered via microinjection into the heart an hour before imaging (4.6nl, 0.5mg/ml); alternatively via mRNA microinjection at the single cell stage (2.3nl, 15ng/*μ*l). This method of anaesthetizing produces fewer developmental delays and defects than the conventional method, tricaine (Swinburne et al., 2015). For endpoint images, in which embryo health was less critical, we used tricaine (Sigma), at 0.2mg/ml.

Prior to imaging, healthy embryos were selected and dechorionated on a glass dish then transferred to a 1.5% agarose 0.4*μ*m canyon mount (Megason, 2009). Using a stereoscope, embryos were carefully positioned within the canyon by a hair loop, with the dorsal neural tube oriented upwards. For the majority of experiments, embryos were mounted in egg water, except some of the embryos for the results in Fig. 2, where they were mounted in 1% low melt agarose (A9414 SIGMA) for increased stability and longer-term imaging.

A Corning coverslip æ1 was placed on top of the agarose mount, taking care not to disturb the embryo positioning.

Imaging was peformed using a Zeiss 710 confocal microscope, C-Apochromat 40X 1.2 NA objective, with a custom made heating chamber to keep the embryos at 28°C. The following laser lines were used: 405nm (eBFP2), 488nm (eGFP), 514nm (citrine) and 594nm (mCherry2). Other parameters were optimized for each experiment (for example, low laser powers were used for all timelapse imaging to prevent bleaching), but were consistent between experimental conditions. Timelapse movies were started at 24hpf ( ± 1hr). Endpoint measurements (Fig. 3C and 5B) were taken at 32hpf.

Figures are composed of single XY slices, dorsal view, of single timepoints from the timelapse data. Note that some images are flipped left to right for consistency of data presentation. The imaging from Fig. S4 was performed on a Nikon Eclipse E800 confocal microscope with the embryos anaesthetized in Tricaine (Sigma) and embedded in low melt agarose (1%) within glass bottomed petri dishes.

### Analysis of timelapse data

Raw Zeiss.lsm files were converted to formats compatible with GoFigure2, an open-source software package to manually analyze *in toto* timelapse imaging data. First, 3-4 cells were manually tracked for the entire length of the movie. These tracks were then used to register the data between timepoints, thus removing global translation and rotation of the embryo. Then, using this registered dataset, we assembled a set of tracks. We started each track at its division (evident by its spherical morphology), and tracked both forwards and backwards in time. We restricted our tracks to cells within the central 30% of the neural tube along DV, and rejected cells that could not be tracked for long periods e.g. those that moved out of the field of view too quickly, or had poor membrane signal. GFP intensity (from *neurod:eGFP*) was used to identify neurons. To positively identify a neuron, we required that the entire cell was GFP positive, in each of the XY, XZ and YZ image planes, to avoid the potential of GFP scatter from neighboring cells giving false positives. The first time at which a cell was identified as a neuron was recorded. In some cases, GFP was excited intermittently throughout the timelapse to reduce bleaching (e.g. every hour, instead of every 3 minutes). In this case, the time recorded was chosen to be midway between the time intervals (e.g. if a cell was negative at 3hrs, but positive at 4hrs, we record 3.5hrs). Cells that did not turn on GFP were tracked either until they divided, or they were no longer trackable, and the total track time was recorded. GFP-on times (‘events’) were combined with the total track time to generate Kaplan-Meier plots (MATLAB). These are commonly used to analyze survival times in the medical community. For example, an ‘event’ could be recovery from a certain illness, and the Kaplan-Meier plots are used to compare recovery time between placebo and drug-treated subjects. They are particularly useful when not all subjects complete the entire study, termed ‘censoring’, as well as for analyzing *in toto* image tracks, which are of variable lengths.

### Manual image analysis

Distances are measured within GoFigure2 using a 3D distance tool. The dorsal-ventral height is measured from the base of the floorplate to the top of the roofplate. The mediolateral width is measured at the point along DV where it is widest. The anterior-posterior segment length is found by measuring the AP distance between neurons that first project ventrally, which occur once per neural hemisegment. Cell (or nuclei) centroid positions are manually identified and recorded by the placement of a cell mesh, and its distance to the apical surface is measured again using the 3D distance tool.

Notch activity was measured by GFP intensity from the nort:dGFP reporter - GFP intensity was measured within a 3*μ*m radius spherical mesh, whose center was placed 12*μ*m away from the apical surface in line with the arrested mitotic cell (BFP positive). For control, two random numbers were chosen (MATLAB) to generate positions along the DV and AP axes, and a 3*μ*m mesh was placed 12*μ*m away from this point.

### Automated image analysis: high quality single timepoint images

Raw Zeiss .lsm files were first converted to .mha files. Segmentation was then performed on the membrane channel, using the ACME algorithm (Mosaliganti et al., 2012). A mask was created in GoFigure2 to correctly identify meshes that fell within the neural tube, and excluded skin, notochord and somite cells. Cell position, volume, shape and median GFP intensity were extracted from the cell meshes, and analyzed in MATLAB. Progenitor density was calculated by binning cells along the DV and AP axes into 14*μ*m bins and counting cell number within each bin. Segmentations and neuron classifications were visually inspected on ITKsnap and, where necessary, manually corrected. All image analysis was performed using custom C++ scripts.

### DNA constructs

The hsp:mTagBFP2-dnPLK1 construct was generated by fusing the coding sequence for mTagBFP2 (gift from Pamela Silver) to a dominant negative human polo-kinase 1 (gift from Caren Norden (Strzyz et al., 2015)), using a flexible GA linker, and inserting into a vector containing the hsp70 promoter (Xiong et al., 2015). The hsp:mTagBFP2-dnPLK1-2A-p50 was similarly made, but with two extra pieces: the P2A sequence (gift from Tony Tsai, Addgene æ52421) and the p50 (amplified from zebrafish cDNA). Pieces were amplified using PCR with 20-30bp overlap regions, and combined using isothermal assembly. RT-PCR was performed to generate TOPO^®^ (Life Technologies) plasmids of the full-length cDNA sequence for zebrafish *nort*. The TOPO® (Life Technologies) *nort* plasmid was used to generate an in situ primer. A list of primers is provided in Table S1.

### Fluorescent in situ hybridization

Dechorionated embryos were fixed in fresh, ice cold 4% paraformaldehyde/PBS overnight at 4°C. After fixation embyros were washed twice in ice cold PBS and then four times in ice cold 100% MeOH and stored in MeOH (15-20 embryos per tube) at -20°C. Following methanol fixation embryos were re-hydrated in a dilution series of MeOH:1xPBS/1%Tween-20 (3:1,1:1,1:3) and then standard in situ methodology (Thisse and Thisse, 2004) was followed and Fast Red tablets (*F4648* Sigma) were used to visualize *nort* mRNA.

### Whole Mount Zebrafish Larvae Immunofluoresence

Embryos were fixed in 4% paraformaldehyde in 1xPBS (pH 7.4) overnight at 4°C and after fixation rinsed 3 times for 5 minutes in 1xPBS. Prior to immunostaining, embyros were blocked for 60 minutes at room temperature in 2% normal goat serum/1%Triton X-100/1% Tween-20/1xPBS (pH 7.4) (blocking buffer). After block, embryos were incubated in diluted anti-Myc primary antibody (clone9E10, Thermofisher) at 1:200 dilution in blocking buffer overnight at room temperature. Embryos were then rinsed in 1% Tween-20/1xPBS (pH 7.4) and then washed 3 times for 60 minutes in 1% Tween-20/1xPBS (pH 7.4). Then embryos were incubated in Alexa-567 secondary antibodies (Invitrogen) at 1:800 in blocking buffer overnight at 4°C. Following secondary treatment embryos were washed 4 times for 30 minutes in 1% Tween-20/1xPBS

### Microinjections of DNA and mRNA

Plasmid DNA (hsp:mTagBFP2-dnPLK1, hsp:mTagBFP2-dnPLK1-P2A-p50) was injected at the single cell stage, delivering 2.3nl (Nanoject) at a concentration of 10ng/*μ*l combined with 25ng/*μ*l transposase mRNA. mRNA (mem-citrine-citrine) was synthesized using the mMESSAGE mMACHINE kits (Ambion), and injected at the 16-128 cell stage for mosaic labeling at a concentration of 20ng/*μ*l. Prior to each experiment, embryos were screened for health. 4.6nl of 10 ng/*μ*l *5xUASE1b*:6xMYC-notch1a (Scheer and Campos-Ortega, 1999) plasmid DNA was injected into 1-4 cell stage embryos.

### Morpholinos (MO)

*tp53* MO, 5**’**-GCGCCATTGCTTTGCAAGAATTG-3 **‘** (Robu et al., 2007). Injected 9.2 nL of a 50 *μ*M MO concentration.

*rbpj* ATG MO 5’ – CAAACTTCCCTGTCACAACAGGCGC – 3’ (Ohata et al., 2011) Injected 9.2 nL of a 50 *μ*M MO concentration.

### Heatshock treatment

Embryos were placed in a 1.5ml Eppendorf tube containing (pre-warmed) egg water, at 37**°**C, for 45 minutes, between 19-20hpf. Embryos were then removed and placed in fresh (22-28**°**C) water, and returned to the incubator.

### Statistical tests

Statistical analysis of pairwise comparisons was mainly performed using a two-tailed t-test (ttest2 in MATLAB). Several variables had a highly skewed distribution (namely: (1) distance of cell to apical surface, and (2) nort:dGFP expression) and so in these cases we used a Mann-Whitney test (ranksum in MATLAB). Kaplan-Meier curves were analyzed using the log rank test (logrank in MATLAB) (Rich et al., 2010). For Figure 2C, we fit the Kaplan-Meier curves to a parametric form *ρ*(*t*) = 1 − exp [*R(t_r_ − t)*] for the first 6 hours after division. Note that *R* can be related to the division probability in Figure 6, provided one knows when cells stop differentiating (e.g. when they enter S phase). The fitting was implemented as a linear fit of ln (1 − ρ (t)) in time. For the plot of *R* as a function of *d*, *R(d)* corresponds to the differentiation rate for all cells whose nuclear distance exceeds the value *d.*

### Mathematical model

A stochastic simulation was implemented in MATLAB. Each progenitor is modeled independently and after birth is assumed to divide again with a cell cycle time taken from the distribution in Fig. 6C (a generalized extreme value distribution (Bogdan et al., 2014)). Upon dividing, each daughter cell differentiates independently with probability *f(t)*. In our simulations, *f(t)* increases linearly from zero to one over the course of 12 cell cycles. For Fig. 6D,E, we repeat each simulation 3000 times and plot the mean, plus/minus the standard deviation as shown. For feedback, we modify *f(t)* → *f(t)* + *κP* − δ where *κ* is a constant controlling the strength of feedback, and κ is a constant that is manually tuned such that the mean dynamics are similar to the case without feedback.

## Supplementary Figure Captions

**Figure S1.**
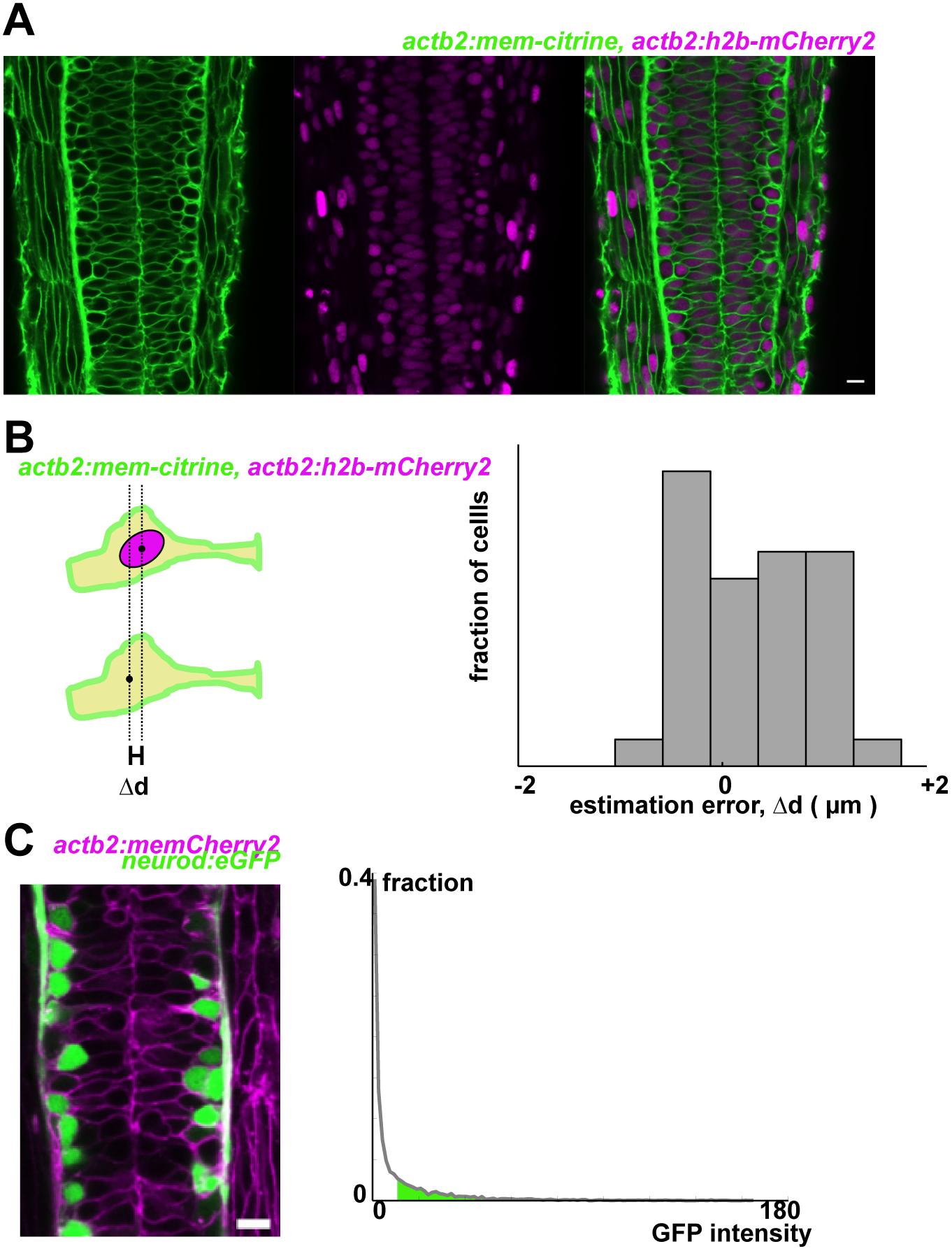
A: Embryos doubly transgenic for a membrane and nuclear label (mem-citrine and h2b-mCherry2) reveal the densely-packed pseudostratified epithelial character of the neural tube. B: We compared nuclear position (based on h2b signal) with the cell centroid position (based on the citrine signal), both manually identified. We find that the difference between these two measurements is rather small (mean value < 1*μ*m). C: *Tg(neurod:eGFP)* is used to classify neurons versus progenitors. Cells are segmented, and the median GFP intensity is calculated per cell (the median, rather than the mean, is robust to scatter of GFP signal from high intensity neighboring cells). Neurons are identified as having a median GFP intensity higher than a certain threshold, defined manually by referencing the raw images, and is fixed for all samples for the same experiment.

**Figure S2.**
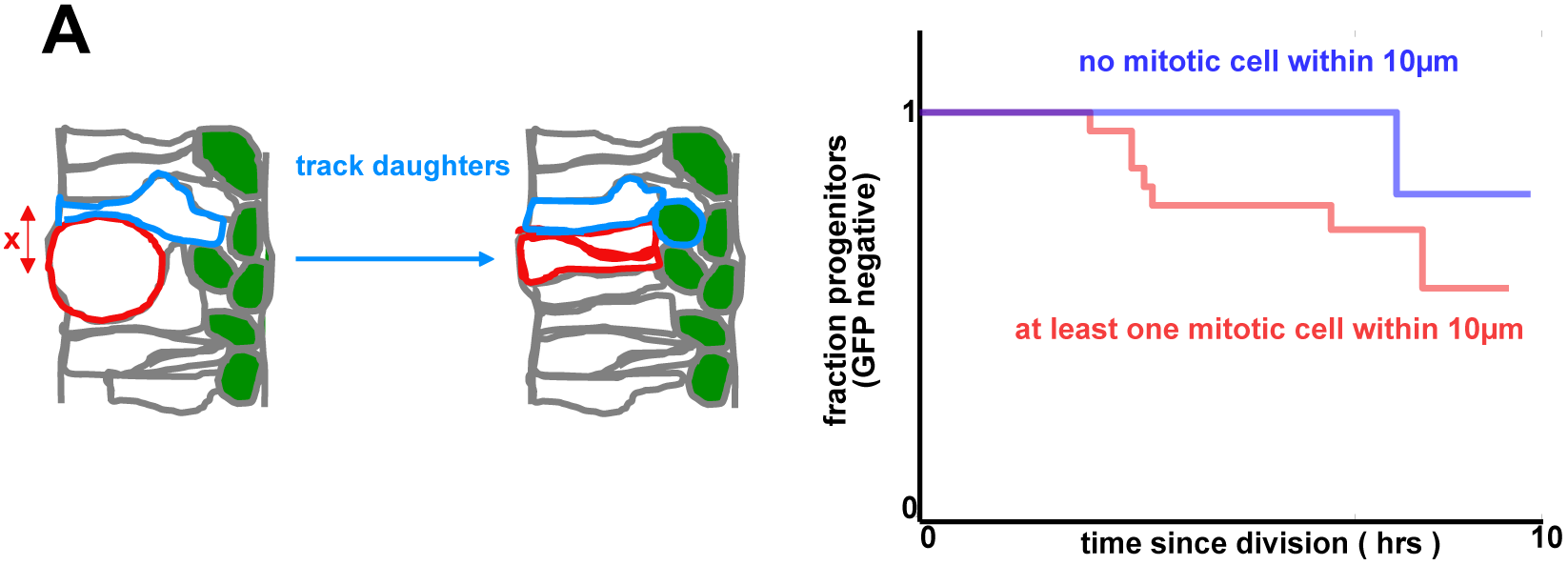
A: By tracking progenitors, we see a small, but not significant (p = 0.1) difference in differentiation rate when comparing cells that are or are not adjacent to dividing cells (n=58, same movies as in Figure 2).

**Figure S3.**
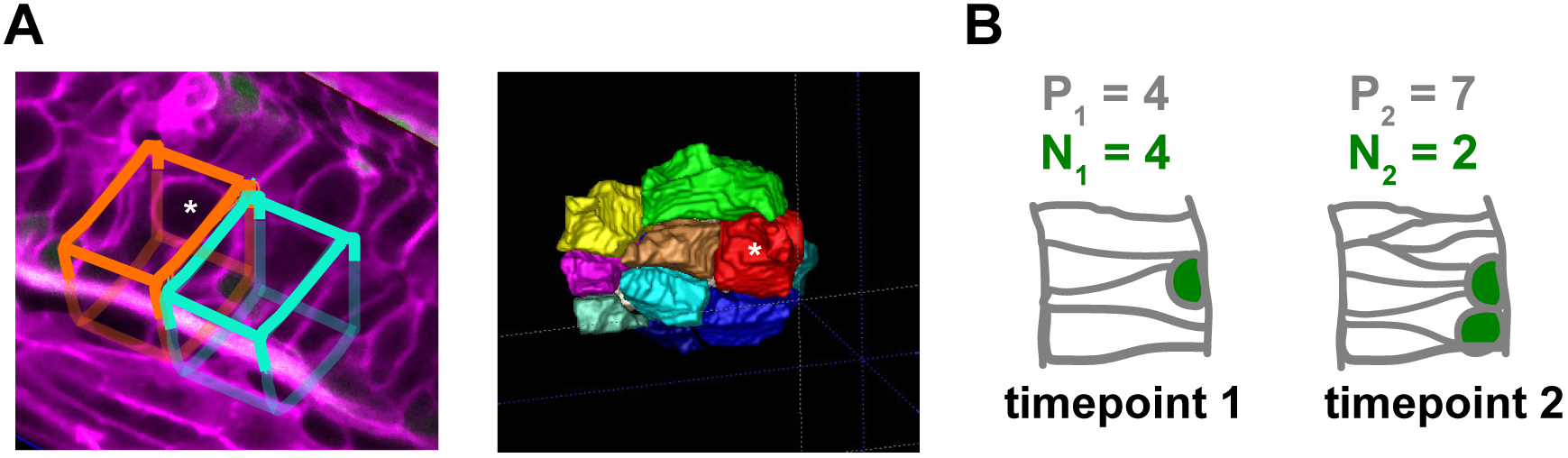
A: Left: We compare regions adjacent to a mitotic cell (15*μ*m x 15*μ*m apical surface, manually contoured) [orange], to nearby control regions of the same dimensions but without an arrested cell [cyan]. Single cell tracking reveals that there is little movement of cells along AP/DV that would take them out of the regions (∼2.6*μ*m mean distance moved). Right: segmented image. Asterisks depict an arrested mitotic cell. B: Illustrative calculation. We define the fraction of neurons as f = N/(N + P), i.e. for the right f = 2/(2 + 7) = 0.22 . We define the proliferation rate between the two timepoints as: μ = (P_2_ + N_2_-P_1_-N_1_)/2(P_1_ + P_2_).

**Figure S4.**
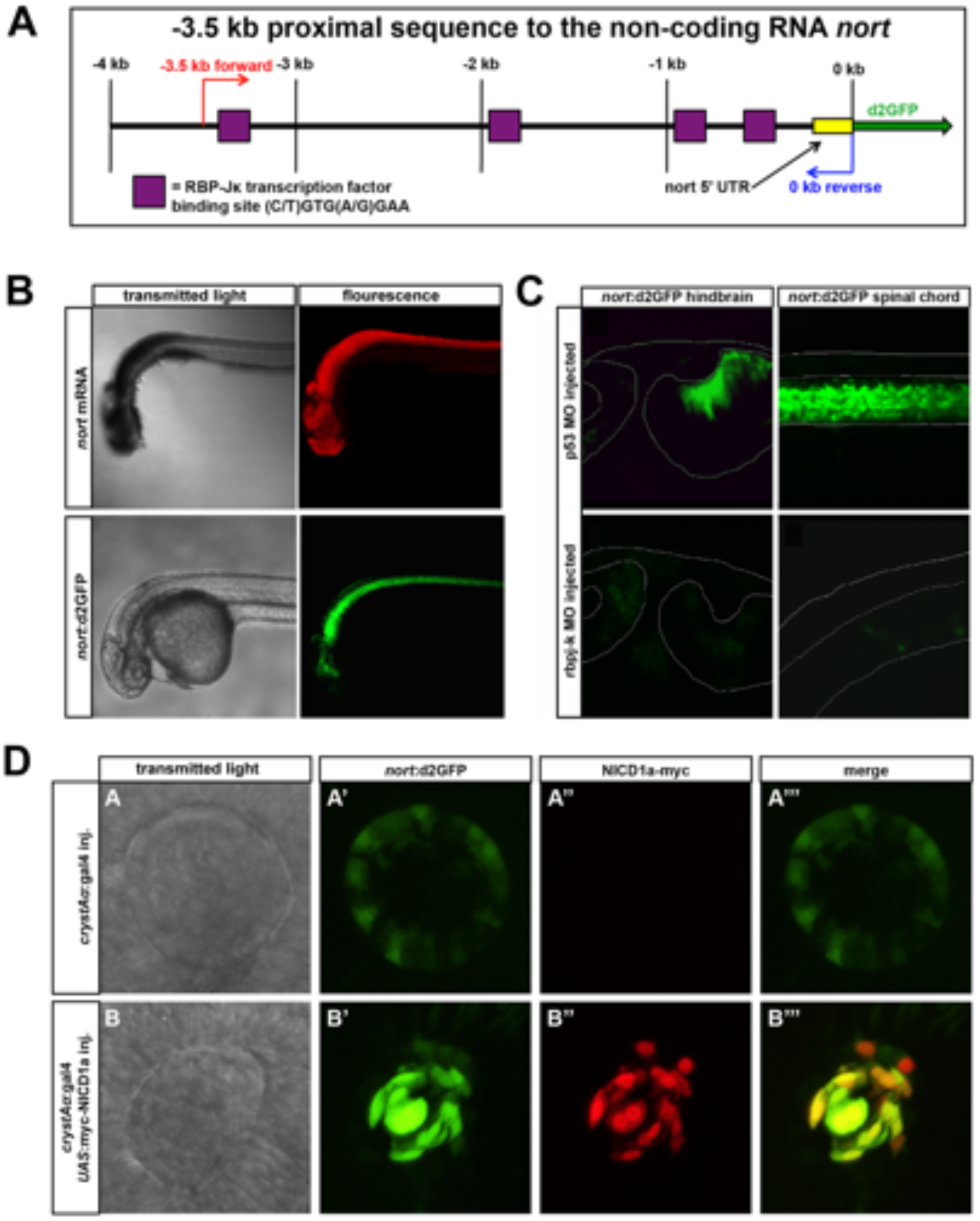
A: Schematic of the 3.5 kb *nort* proximal promoter. This region contains 4 canonical *rbpj-K* binding sites. B: *nort* mRNA (red) and *nort*:d2GFP (green) transgene express in the same tissues at 30 hpf. C: *nort*:d2GFP expression was reduced in a rbpj-k MO injected 48 hpf hindbrain and spinal cord compared to the p53 MO injected controls. The morphology of the hindbrain was similar to the controls while the rbpj-k MO injected spinal cord was curved. D: Normal expression of *nort*:d2GFP is found in the lens epithelium. Overexpression of myc-NICD1a in the lens using *crystA α*:gal4 causes enhanced ectopic *nort*:d2GFP expression. myc immunofluorescence co-localizes with upregulated d2GFP at 28 hpf.

**Table S1:**
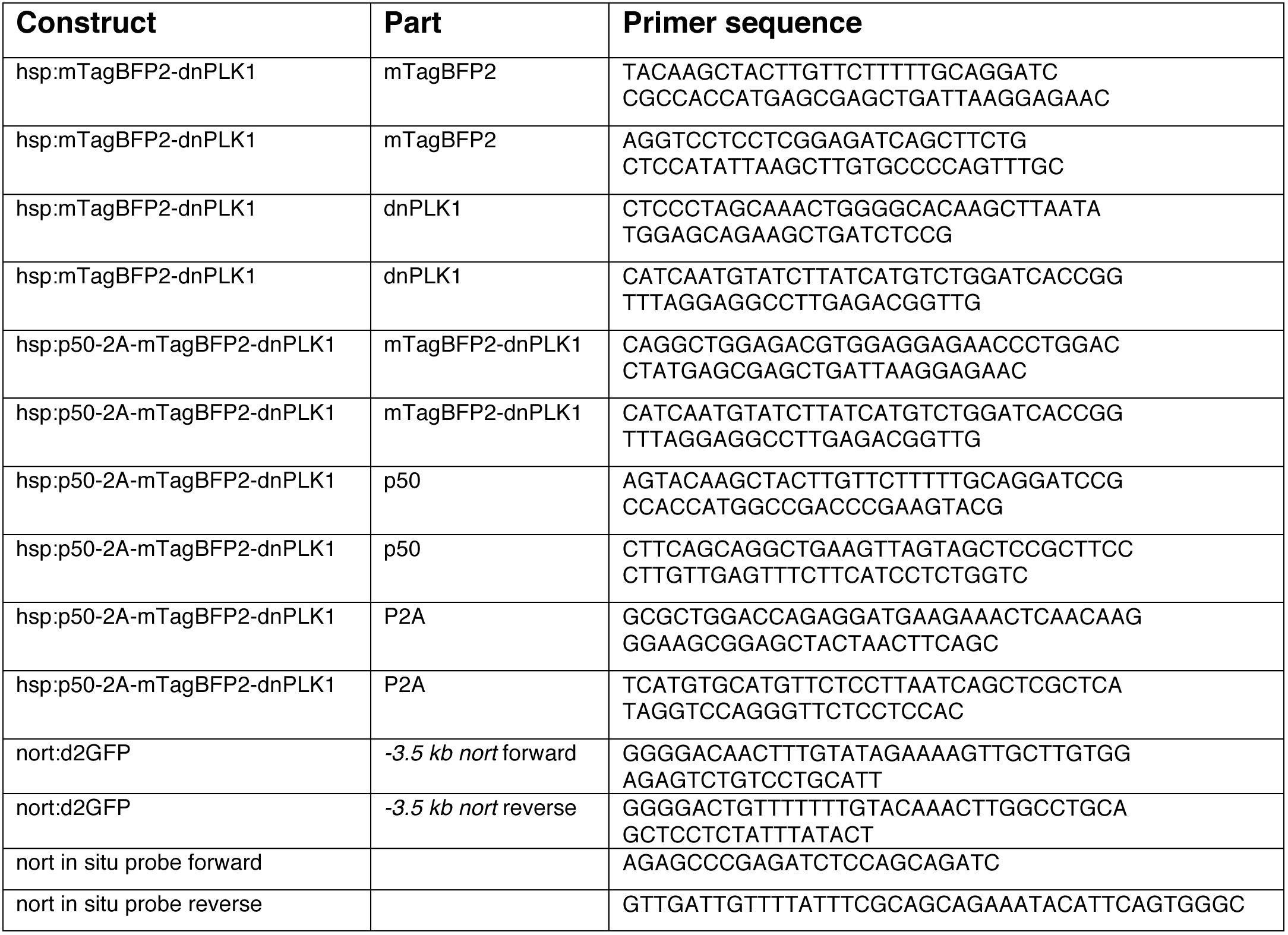
Primers used in this study

## References

Aggarwal, V., Dickinson, R.B., Lele, T.P., 2016. Concentration Sensing by the Moving Nucleus in Cell Fate Determination: A Computational Analysis. PloS one 11, e0149213.

Alexandre, P., Reugels, A.M., Barker, D., Blanc, E., Clarke, J.D., 2010. Neurons derive from the more apical daughter in asymmetric divisions in the zebrafish neural tube. Nature neuroscience 13, 673–679.

Alexiades, M.R., Cepko, C., 1996. Quantitative analysis of proliferation and cell cycle length during development of the rat retina. Developmental dynamics : an official publication of the American Association of Anatomists 205, 293–307.

Appel, B., Givan, L.A., Eisen, J.S., 2001. Delta-Notch signaling and lateral inhibition in zebrafish spinal cord development. BMC developmental biology 1, 13.

Aragona, M., Panciera, T., Manfrin, A., Giulitti, S., Michielin, F., Elvassore, N., Dupont, S., Piccolo, S., 2013. A mechanical checkpoint controls multicellular growth through YAP/TAZ regulation by actin-processing factors. Cell 154, 1047–1059.

Arulmoli, J., Pathak, M.M., McDonnell, L.P., Nourse, J.L., Tombola, F., Earthman, J.C., Flanagan, L.A., 2015. Static stretch affects neural stem cell differentiation in an extracellular matrix-dependent manner. Scientific reports 5, 8499.

Baye, L.M., Link, B.A., 2007. Interkinetic nuclear migration and the selection of neurogenic cell divisions during vertebrate retinogenesis. The Journal of neuroscience : the official journal of the Society for Neuroscience 27, 10143–10152.

Benham-Pyle, B.W., Pruitt, B.L., Nelson, W.J., 2015. Cell adhesion. Mechanical strain induces E-cadherin-dependent Yap1 and beta-catenin activation to drive cell cycle entry. Science 348, 1024–1027.

Bogdan, P., Deasy, B.M., Gharaibeh, B., Roehrs, T., Marculescu, R., 2014. Heterogeneous structure of stem cells dynamics: statistical models and quantitative predictions. Scientific reports 4, 4826.

Bort, R., Signore, M., Tremblay, K., Martinez Barbera, J.P., Zaret, K.S., 2006. Hex homeobox gene controls the transition of the endoderm to a pseudostratified, cell emergent epithelium for liver bud development. Developmental biology 290, 44–56.

Bray, S.J., 2016. Notch signalling in context. Nature reviews. Molecular cell biology 17, 722–735.

Brunet, T., Bouclet, A., Ahmadi, P., Mitrossilis, D., Driquez, B., Brunet, A.C., Henry, L., Serman, F., Bealle, G., Menager, C., Dumas-Bouchiat, F., Givord, D., Yanicostas, C., Le-Roy, D., Dempsey, N.M., Plessis, A., Farge, E., 2013. Evolutionary conservation of early mesoderm specification by mechanotransduction in Bilateria. Nature communications 4, 2821.

Burkhardt, J.K., Echeverri, C.J., Nilsson, T., Vallee, R.B., 1997. Overexpression of the dynamitin (p50) subunit of the dynactin complex disrupts dynein-dependent maintenance of membrane organelle distribution. The Journal of cell biology 139, 469–484.

Clark, B.S., Cui, S., Miesfeld, J.B., Klezovitch, O., Vasioukhin, V., Link, B.A., 2012. Loss of Llgl1 in retinal neuroepithelia reveals links between apical domain size, Notch activity and neurogenesis. Development 139, 1599–1610.

Copp, A.J., Stanier, P., Greene, N.D., 2013. Neural tube defects: recent advances, unsolved questions, and controversies. The Lancet. Neurology 12, 799–810.

Coste, B., Xiao, B., Santos, J.S., Syeda, R., Grandl, J., Spencer, K.S., Kim, S.E., Schmidt, M., Mathur, J., Dubin, A.E., Montal, M., Patapoutian, A., 2012. Piezo proteins are pore-forming subunits of mechanically activated channels. Nature 483, 176–181.

Del Bene, F., Wehman, A.M., Link, B.A., Baier, H., 2008. Regulation of neurogenesis by interkinetic nuclear migration through an apical-basal notch gradient. Cell 134, 1055–1065.

Dessaud, E., Yang, L.L., Hill, K., Cox, B., Ulloa, F., Ribeiro, A., Mynett, A., Novitch, B.G., Briscoe, J., 2007. Interpretation of the sonic hedgehog morphogen gradient by a temporal adaptation mechanism. Nature 450, 717–720.

Dong, Z., Yang, N., Yeo, S.Y., Chitnis, A., Guo, S., 2012. Intralineage directional Notch signaling regulates self-renewal and differentiation of asymmetrically dividing radial glia. Neuron 74, 65–78.

Dupont, S., Morsut, L., Aragona, M., Enzo, E., Giulitti, S., Cordenonsi, M., Zanconato, F., Le Digabel, J., Forcato, M., Bicciato, S., Elvassore, N., Piccolo, S., 2011. Role of YAP/TAZ in mechanotransduction. Nature 474, 179–183.

Engler, A.J., Sen, S., Sweeney, H.L., Discher, D.E., 2006. Matrix elasticity directs stem cell lineage specification. Cell 126, 677–689.

Fernandez-Sanchez, M.E., Barbier, S., Whitehead, J., Bealle, G., Michel, A., Latorre-Ossa, H., Rey, C., Fouassier, L., Claperon, A., Brulle, L., Girard, E., Servant, N., Rio-Frio, T., Marie, H., Lesieur, S., Housset, C., Gennisson, J.L., Tanter, M., Menager, C., Fre, S., Robine, S., Farge, E., 2015. Mechanical induction of the tumorigenic beta-catenin pathway by tumour growth pressure. Nature 523, 92–95.

Garcia-Campmany, L., Marti, E., 2007. The TGFbeta intracellular effector Smad3 regulates neuronal differentiation and cell fate specification in the developing spinal cord. Development 134, 65–75.

Gilbert, P.M., Havenstrite, K.L., Magnusson, K.E., Sacco, A., Leonardi, N.A., Kraft, P., Nguyen, N.K., Thrun, S., Lutolf, M.P., Blau, H.M., 2010. Substrate elasticity regulates skeletal muscle stem cell self-renewal in culture. Science 329, 1078–1081.

Gordon, W.R., Zimmerman, B., He, L., Miles, L.J., Huang, J., Tiyanont, K., McArthur, D.G., Aster, J.C., Perrimon, N., Loparo, J.J., Blacklow, S.C., 2015. Mechanical Allostery: Evidence for a Force Requirement in the Proteolytic Activation of Notch. Developmental cell 33, 729–736.

Grosse, A.S., Pressprich, M.F., Curley, L.B., Hamilton, K.L., Margolis, B., Hildebrand, J.D., Gumucio, D.L., 2011. Cell dynamics in fetal intestinal epithelium: implications for intestinal growth and morphogenesis. Development 138, 4423–4432.

Hardwick, L.J., Ali, F.R., Azzarelli, R., Philpott, A., 2015. Cell cycle regulation of proliferation versus differentiation in the central nervous system. Cell and tissue research 359, 187–200.

Hardwick, L.J., Philpott, A., 2014. Nervous decision-making: to divide or differentiate. Trends in genetics : TIG 30, 254–261.

Hatakeyama, J., Wakamatsu, Y., Nagafuchi, A., Kageyama, R., Shigemoto, R., Shimamura, K., 2014. Cadherin-based adhesions in the apical endfoot are required for active Notch signaling to control neurogenesis in vertebrates. Development 141, 1671–1682.

Hayes, J.M., Hartsock, A., Clark, B.S., Napier, H.R., Link, B.A., Gross, J.M., 2012. Integrin alpha5/fibronectin1 and focal adhesion kinase are required for lens fiber morphogenesis in zebrafish. Molecular biology of the cell 23, 4725–4738.

He, J., Zhang, G., Almeida, A.D., Cayouette, M., Simons, B.D., Harris, W.A., 2012. How variable clones build an invariant retina. Neuron 75, 786–798.

Hindley, C., Philpott, A., 2012. Co-ordination of cell cycle and differentiation in the developing nervous system. The Biochemical journal 444, 375–382.

Huang, P., Xiong, F., Megason, S.G., Schier, A.F., 2012. Attenuation of Notch and Hedgehog signaling is required for fate specification in the spinal cord. PLoS genetics 8, e1002762.

Hudish, L.I., Galati, D.F., Ravanelli, A.M., Pearson, C.G., Huang, P., Appel, B., 2016. miR-219 regulates neural progenitors by dampening apical Par protein-dependent Hedgehog signaling. Development 143, 2292–2304.

Huttner, W.B., Kosodo, Y., 2005. Symmetric versus asymmetric cell division during neurogenesis in the developing vertebrate central nervous system. Current opinion in cell biology 17, 648–657.

Jinguji, Y., Ishikawa, H., 1992. Electron microscopic observations on the maintenance of the tight junction during cell division in the epithelium of the mouse small intestine. Cell structure and function 17, 27–37.

Kicheva, A., Bollenbach, T., Ribeiro, A., Valle, H.P., Lovell-Badge, R., Episkopou, V., Briscoe, J., 2014. Coordination of progenitor specification and growth in mouse and chick spinal cord. Science 345, 1254927.

Kosodo, Y., Suetsugu, T., Suda, M., Mimori-Kiyosue, Y., Toida, K., Baba, S.A., Kimura, A., Matsuzaki, F., 2011. Regulation of interkinetic nuclear migration by cell cycle-coupled active and passive mechanisms in the developing brain. The EMBO journal 30, 1690–1704.

Kwan, K.M., Fujimoto, E., Grabher, C., Mangum, B.D., Hardy, M.E., Campbell, D.S., Parant, J.M., Yost, H.J., Kanki, J.P., Chien, C.B., 2007. The Tol2kit: a multisite gateway-based construction kit for Tol2 transposon transgenesis constructs. Dev Dyn 236, 3088–3099.

Latasa, M.J., Cisneros, E., Frade, J.M., 2009. Cell cycle control of Notch signaling and the functional regionalization of the neuroepithelium during vertebrate neurogenesis. The International journal of developmental biology 53, 895–908.

Le Dreau, G., Saade, M., Gutierrez-Vallejo, I., Marti, E., 2014. The strength of SMAD1/5 activity determines the mode of stem cell division in the developing spinal cord. The Journal of cell biology 204, 591–605.

Lee, J.E., 1997. Basic helix-loop-helix genes in neural development. Current opinion in neurobiology 7, 13–20.

Leipzig, N.D., Shoichet, M.S., 2009. The effect of substrate stiffness on adult neural stem cell behavior. Biomaterials 30, 6867–6878.

Leung, L., Klopper, A.V., Grill, S.W., Harris, W.A., Norden, C., 2011. Apical migration of nuclei during G2 is a prerequisite for all nuclear motion in zebrafish neuroepithelia. Development 138, 5003–5013.

Megason, S.G., 2009. In toto imaging of embryogenesis with confocal time-lapse microscopy. Methods in molecular biology 546, 317–332.

Miguez, D.G., 2015. A Branching Process to Characterize the Dynamics of Stem Cell Differentiation. Scientific reports 5, 13265.

Mosaliganti, K.R., Noche, R.R., Xiong, F., Swinburne, I.A., Megason, S.G., 2012. ACME: automated cell morphology extractor for comprehensive reconstruction of cell membranes. PLoS computational biology 8, e1002780.

Noctor, S.C., Martinez-Cerdeno, V., Ivic, L., Kriegstein, A.R., 2004. Cortical neurons arise in symmetric and asymmetric division zones and migrate through specific phases. Nature neuroscience 7, 136–144.

Norden, C., Young, S., Link, B.A., Harris, W.A., 2009. Actomyosin is the main driver of interkinetic nuclear migration in the retina. Cell 138, 1195–1208.

Nowell, C.S., Odermatt, P.D., Azzolin, L., Hohnel, S., Wagner, E.F., Fantner, G.E., Lutolf, M.P., Barrandon, Y., Piccolo, S., Radtke, F., 2016. Chronic inflammation imposes aberrant cell fate in regenerating epithelia through mechanotransduction. Nature cell biology 18, 168–180.

Obholzer, N., Wolfson, S., Trapani, J.G., Mo, W., Nechiporuk, A., Busch-Nentwich, E., Seiler, C., Sidi, S., Sollner, C., Duncan, R.N., Boehland, A., Nicolson, T., 2008. Vesicular glutamate transporter 3 is required for synaptic transmission in zebrafish hair cells. The Journal of neuroscience : the official journal of the Society for Neuroscience 28, 2110–2118.

Ohata, S., Aoki, R., Kinoshita, S., Yamaguchi, M., Tsuruoka-Kinoshita, S., Tanaka, H., Wada, H., Watabe, S., Tsuboi, T., Masai, I., Okamoto, H., 2011. Dual roles of Notch in regulation of apically restricted mitosis and apicobasal polarity of neuroepithelial cells. Neuron 69, 215–230.

Otani, T., Marchetto, M.C., Gage, F.H., Simons, B.D., Livesey, F.J., 2016. 2D and 3D Stem Cell Models of Primate Cortical Development Identify Species-Specific Differences in Progenitor Behavior Contributing to Brain Size. Cell stem cell 18, 467–480.

Ou, G., Weaver, V.M., 2015. Tumor-induced solid stress activates beta-catenin signaling to drive malignant behavior in normal, tumor-adjacent cells. BioEssays : news and reviews in molecular, cellular and developmental biology 37, 1293–1297.

Pan, Y., Heemskerk, I., Ibar, C., Shraiman, B.I., Irvine, K.D., 2016. Differential growth triggers mechanical feedback that elevates Hippo signaling. Proceedings of the National Academy of Sciences of the United States of America.

Paolini, A., Duchemin, A.L., Albadri, S., Patzel, E., Bornhorst, D., Gonzalez Avalos, P., Lemke, S., Machate, A., Brand, M., Sel, S., Di Donato, V., Del Bene, F., Zolessi, F.R., Ramialison, M., Poggi, L., 2015. Asymmetric inheritance of the apical domain and self-renewal of retinal ganglion cell progenitors depend on Anillin function. Development 142, 832–839.

Rich, J.T., Neely, J.G., Paniello, R.C., Voelker, C.C., Nussenbaum, B., Wang, E.W., 2010. A practical guide to understanding Kaplan-Meier curves. Otolaryngology--head and neck surgery : official journal of American Academy of Otolaryngology-Head and Neck Surgery 143, 331–336.

Robu, M.E., Larson, J.D., Nasevicius, A., Beiraghi, S., Brenner, C., Farber, S.A., Ekker, S.C., 2007. p53 activation by knockdown technologies. PLoS Genet 3, e78.

Saade, M., Gutierrez-Vallejo, I., Le Dreau, G., Rabadan, M.A., Miguez, D.G., Buceta, J., Marti, E., 2013. Sonic hedgehog signaling switches the mode of division in the developing nervous system. Cell reports 4, 492–503.

Scheer, N., Campos-Ortega, J.A., 1999. Use of the Gal4-UAS technique for targeted gene expression in the zebrafish. Mechanisms of development 80, 153–158.

Schier, A.F., Neuhauss, S.C., Harvey, M., Malicki, J., Solnica-Krezel, L., Stainier, D.Y., Zwartkruis, F., Abdelilah, S., Stemple, D.L., Rangini, Z., Yang, H., Driever, W., 1996. Mutations affecting the development of the embryonic zebrafish brain. Development 123, 165–178.

Seidlits, S.K., Khaing, Z.Z., Petersen, R.R., Nickels, J.D., Vanscoy, J.E., Shear, J.B., Schmidt, C.E., 2010. The effects of hyaluronic acid hydrogels with tunable mechanical properties on neural progenitor cell differentiation. Biomaterials 31, 3930–3940.

Shaya, O., Binshtok, U., Hersch, M., Rivkin, D., Weinreb, S., Amir-Zilberstein, L., Khamaisi, B., Oppenheim, O., Desai, R.A., Goodyear, R.J., Richardson, G.P., Chen, C.S., Sprinzak, D., 2017. Cell-Cell Contact Area Affects Notch Signaling and Notch-Dependent Patterning. Developmental cell 40, 505–511 e506.

Shraiman, B.I., 2005. Mechanical feedback as a possible regulator of tissue growth. Proceedings of the National Academy of Sciences of the United States of America 102, 3318–3323.

Spear, P.C., Erickson, C.A., 2012. Interkinetic nuclear migration: a mysterious process in search of a function. Development, growth & differentiation 54, 306–316.

Streichan, S.J., Hoerner, C.R., Schneidt, T., Holzer, D., Hufnagel, L., 2014. Spatial constraints control cell proliferation in tissues. Proceedings of the National Academy of Sciences of the United States of America 111, 5586–5591.

Strzyz, P.J., Lee, H.O., Sidhaye, J., Weber, I.P., Leung, L.C., Norden, C., 2015. Interkinetic nuclear migration is centrosome independent and ensures apical cell division to maintain tissue integrity. Developmental cell 32, 203–219.

Swinburne, I.A., Mosaliganti, K.R., Green, A.A., Megason, S.G., 2015. Improved Long-Term Imaging of Embryos with Genetically Encoded alpha-Bungarotoxin. PloS one 10, e0134005.

Tallinen, T., Chung, J.Y., Biggins, J.S., Mahadevan, L., 2014. Gyrification from constrained cortical expansion. Proceedings of the National Academy of Sciences of the United States of America 111, 12667–12672.

Tsuda, S., Kitagawa, T., Takashima, S., Asakawa, S., Shimizu, N., Mitani, H., Shima, A., Tsutsumi, M., Hori, H., Naruse, K., Ishikawa, Y., Takeda, H., 2010. FAK-mediated extracellular signals are essential for interkinetic nuclear migration and planar divisions in the neuroepithelium. Journal of cell science 123, 484–496.

Tsujikawa, M., Omori, Y., Biyanwila, J., Malicki, J., 2007. Mechanism of positioning the cell nucleus in vertebrate photoreceptors. Proceedings of the National Academy of Sciences of the United States of America 104, 14819–14824.

Tsutsumi, M., Itoh, M., 2007. Novel transcript nort is a downstream target gene of the Notch signaling pathway in zebrafish. Gene expression patterns : GEP 7, 227–232.

Xiong, F., Ma, W., Hiscock, T.W., Mosaliganti, K.R., Tentner, A.R., Brakke, K.A., Rannou, N., Gelas, A., Souhait, L., Swinburne, I.A., Obholzer, N.D., Megason, S.G., 2014. Interplay of cell shape and division orientation promotes robust morphogenesis of developing epithelia. Cell 159, 415–427.

Xiong, F., Obholzer, N.D., Noche, R.R., Megason, S.G., 2015. Multibow: digital spectral barcodes for cell tracing. PloS one 10, e0127822.

Xiong, F., Tentner, A.R., Huang, P., Gelas, A., Mosaliganti, K.R., Souhait, L., Rannou, N., Swinburne, I.A., Obholzer, N.D., Cowgill, P.D., Schier, A.F., Megason, S.G., 2013. Specified neural progenitors sort to form sharp domains after noisy Shh signaling. Cell 153, 550–561.

Yeo, S.Y., Chitnis, A.B., 2007. Jagged-mediated Notch signaling maintains proliferating neural progenitors and regulates cell diversity in the ventral spinal cord. Proceedings of the National Academy of Sciences of the United States of America 104, 5913–5918.

You, H., Sim, K.B., Wang, K.C., Kim, D.G., Kim, H.J., 1994. Morphological study of surgically induced open neural tube defects in chick embryos--postoperative 24 hours. Journal of Korean medical science 9, 116–122.

Zechner, D., Fujita, Y., Hulsken, J., Muller, T., Walther, I., Taketo, M.M., Crenshaw, E.B., 3rd, Birchmeier, W., Birchmeier, C., 2003. beta-Catenin signals regulate cell growth and the balance between progenitor cell expansion and differentiation in the nervous system. Developmental biology 258, 406–418.

